# A Dual-sensing DNA Nanostructure with an Ultra-broad Detection Range

**DOI:** 10.1101/727610

**Authors:** Byunghwa Kang, Soyeon V. Park, H. Tom Soh, Seung Soo Oh

**Affiliations:** Department of Materials Science and Engineering, Pohang University of Science and Technology (POSTECH), 77 Cheongam-Ro, Nam-Gu, Pohang, Gyeongbuk 37673, South Korea; Department of Electrical Engineering and Department of Radiology, Canary Center at Stanford University, 3155 Porter Drive, Stanford, CA 94305 (USA); School of Interdisciplinary Bioscience and Bioengineering, Pohang University of Science and Technology (POSTECH), 77 Cheongam-Ro, Nam-Gu, Pohang, Gyeongbuk 37673, South Korea

## Abstract

Despite considerable interest in the development of biosensors that can measure analyte concentrations with a dynamic range spanning many orders of magnitude, this goal has proven difficult to achieve. We describe here a modular biosensor architecture that integrates two different readout mechanisms into a single-molecule construct that can achieve target detection across an extraordinarily broad dynamic range. Our dual-mode readout DNA biosensor (DMRD) combines an aptamer and a DNAzyme to quantify ATP with two different mechanisms, which respond to low (micromolar) and high (millimolar) concentrations by generating distinct readouts based on changes in fluorescence and absorbance, respectively. Importantly, we have also devised regulatory strategies to finely tune the target detection range of each sensor module by controlling the target-sensitivity of each readout mechanism. Using this strategy, we report the detection of ATP at a dynamic range spanning 1–500,000 μM—more than five orders of magnitude, representing the largest dynamic range reported to date with a single biosensor construct.

## Introduction

In nature, many biological receptors have evolved the capacity to respond to ligands across a broad dynamic range of concentrations.^1–2^ This is important, as their cognate molecular targets, such as hormones, metal ions, and viruses, may be present across a broad spectrum of concentrations.^3–5^ For instance, the JAK2/STAT5 signaling pathway, which is involved in cell differentiation and survival, is capable of responding to a 1,000-fold change in the concentration of the hormone erythropoietin (Epo).^6^ This pathway achieves the broad dynamic range for Epo response through the combined action of two regulatory mechanisms, governed by CIS and SOCS3.^6^ In this signaling pathway, binding of Epo to its receptor induces JAK2 phosphorylation, followed by the activation of the transcription factor STAT5; STAT5 subsequently binds to the Epo receptor to generate a signaling output.^7^ At low levels of Epo (10^−9^ U/cell), CIS acts as a physical competitor that blocks STAT5 binding to the Epo receptor; at higher Epo levels (10^−6^ U/cell), SOCS3 acts as a catalytic inhibitor for Epo-induced JAK2 phosphorylation. The combined action of these two regulators enables cells to respond to changes in Epo concentrations spanning a 1,000-fold range.

Unfortunately, most synthetic biosensors have a much narrower detection range. For example, typical biosensors based on affinity reagents such as antibodies rely on Langmuir binding physics to detect targets, such that their dynamic ranges are both inherently narrow (*i.e*. typically on the order of 81-fold) and extremely difficult to shift.^8–9^ Enzyme-based detection systems likewise have narrowly defined detection ranges because of intrinsic features of the enzyme in question, limiting control over the sensor dynamic range.^10^ These fundamental limitations make it difficult to cope with wide variations in target concentration.^11–12^ Many researchers have proposed ideas to extend the dynamic range of biosensors. For example, Ricci *et al.* have designed diverse DNA-based sensors that can achieve target detection across three orders of magnitude by making use of mutation- or allosteric regulation-based approaches.^13–16^ However, even broader detection ranges are needed for many clinically relevant biological targets; for example, the physiological concentrations of HIV and prostate-specific antigen can vary over more than four orders of magnitude.^5, 17–18^ Therefore, there remains a strong need for novel strategies for the development of simple molecular biosensors with ultra-broad detection ranges.

We present here a novel biosensor design that couples together two DNA-based modules with distinct sensing modalities, enabling semi-quantitative target detection over a concentration range spanning more than five orders of magnitude. Our ‘dual-mode readout DNA’ (DMRD) nanostructure is capable of detecting ATP at two different concentration ranges with two different readouts: fluorescence and absorbance. Moreover, the target detection range of the DMRD is highly tunable, and we have developed strategies for refining the sensitivity of both modules to shape the concentration range at which the sensor produces a response. Using this tuning approach, we demonstrate that our DMRD can successfully report ATP concentrations ranging from 1 μM to 500 mM—a dynamic range spanning a 500,000-fold concentration difference that is presently impossible to obtain with either sensing mechanism alone, and represents the widest target detection range reported to date for a single DNA-based biosensor construct.^13^

## Results and Discussion

### The modular design of the DMRD

As a demonstration of our DMRD concept, we developed a modular ATP sensor (**Figure 1A**). The DMRD comprises two different sensing modules: a direct detection module (DDM), and an enzymatic detection module (EDM). The DDM (**Figure 1A**, top) is an adaptation of a previously published ATP aptamer, which has been engineered to fluoresce in a concentration-dependent manner.^19^ The DDM consists of two single-stranded DNA fragments that have been respectively labeled with fluorescein (FAM) and Black Hole Quencher 1 (BHQ1). These strands remain separated in the absence of ATP, generating an unquenched fluorescent signal. When ATP is added, these two fragments undergo binding-induced assembly that brings FAM and BHQ1 into close proximity, leading to a decrease of fluorescent signal at 535 nm. This aptamer-based DDM is designed to detect relatively low ATP concentrations (≥1 μM).

**Figure 1.**
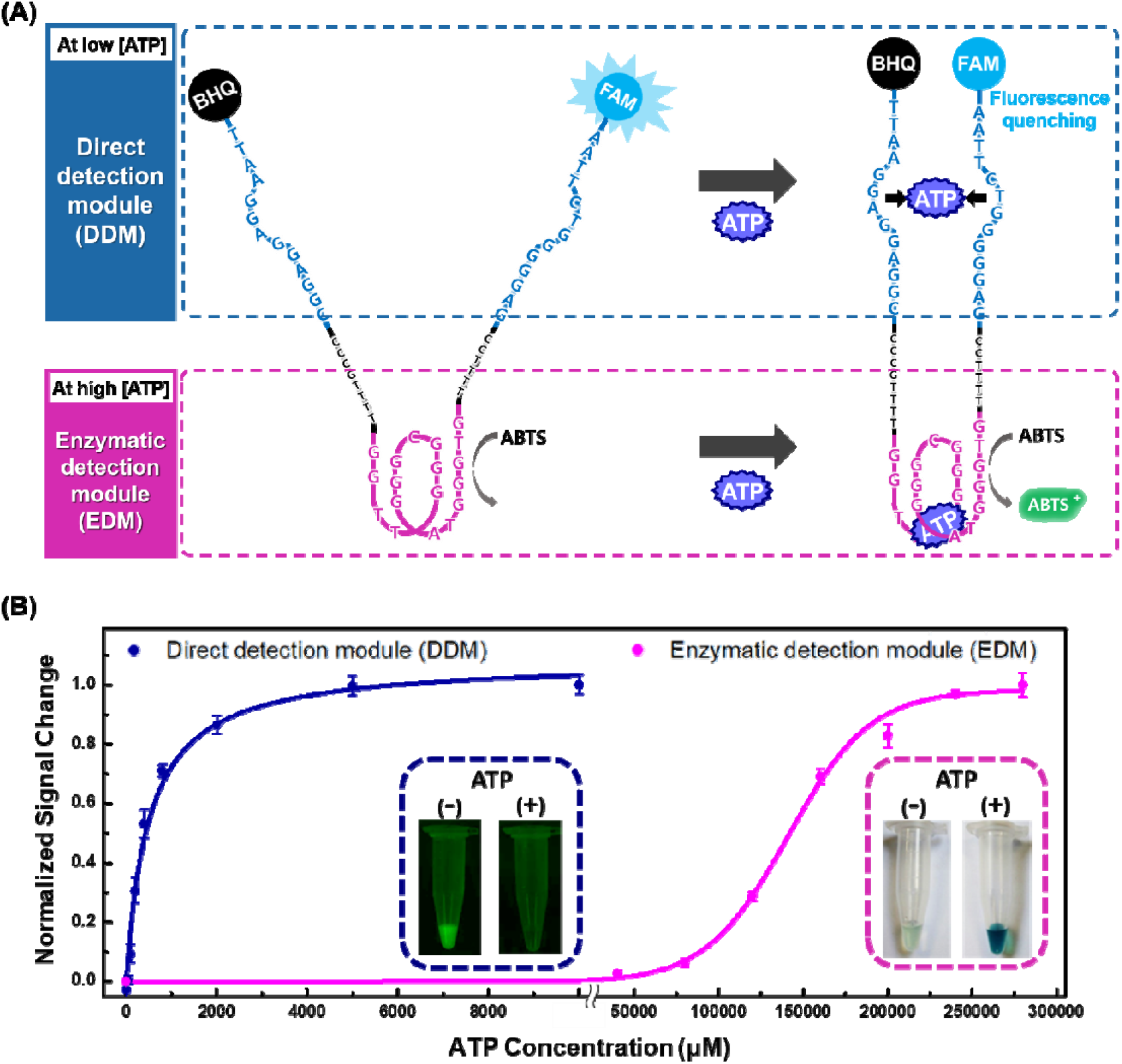
A modular ATP-sensing platform that can achieve a broad dynamic range of detection. (A) Our dual-mode readout DNA (DMRD) nanostructure contains two distinct ATP-sensing modules. The direct detection module (DDM) uses a split ATP-binding aptamer (top), where target binding induces aptamer assembly and thereby brings coupled fluorophore and quencher groups into close proximity. This leads to a measurable decrease in FAM signal that correlates with ATP levels at relatively low (micromolar) concentrations. The enzymatic detection module (EDM) comprises an ATP-enhanced DNAzyme that uses ATP as a cofactor (bottom) that increases its catalytic activity, leading to increased oxidation of ABTS. The accumulation of ABTS^+^ can be measured by a change in absorbance, enabling detection of ATP at higher (millimolar) concentrations. (B) Target detection with the DMRD. The DDM detects ATP in the micromolar concentration range (blue line), and we can observe the ATP-induced decrease in FAM fluorescence at 535 nm (blue inset). The peroxidase activity of the EDM is accelerated by the presence of ATP at millimolar concentrations (magenta line), and ABTS^+^ accumulation leads to increased absorbance at 418 nm that can be observed by naked eye (magenta inset).

The two fragments of the DDM are in turn linked to the ends of the EDM, which provides the second readout mechanism (**Figure 1A**, bottom). The EDM incorporates a DNAzyme, which folds into a G-quadruplex structure that forms a complex with hemin.^20^ This DNAzyme/hemin complex exhibits modest peroxidase activity, but this catalytic activity is greatly enhanced by the presence of ATP.^21–22^ Such ATP-assisted peroxidation can cause rapid oxidation of substrates such as 2,2’-azino-bis(3-ethylbenzothiazoline-6-sulphonic acid) (ABTS), leading to accumulation of the product ABTS^+^. This ABTS^+^ accumulation produces a proportional absorbance change at 418 nm,^23^ generating a colorimetric signal that can be detected by the naked eye at high ATP concentrations (>1 mM).

These two modules can thus be combined to measure the presence of ATP at very different concentration ranges (**Figure 1B**). The DDM, which has a dissociation constant (K_d_) of 23.1 μM, has the capacity to report ATP concentrations in the range of ~1–100 μM based on fluorescence measurements. On the other hand, the EDM exhibits target-assisted peroxidase activity in a millimolar concentration range spanning roughly 0.74–2.92 mM, with the ATP-dependent absorbance change producing a strong green color that is even detectable by naked eye. DDM sensing has no impact on the peroxidase activity of the EDM, which only produces a response at millimolar ATP concentrations at which the DDM’s fluorescent signal has already become saturated. As a consequence, our modular DMRD design can cover a broad dynamic range for target detection through its dual-readout system.

### Optimizing DDM performance

The detection range of the DDM can be finely tuned through the use of antisense DNAs that modulate aptamer-target binding (**Figure 2A**), yielding the capacity to discriminate ATP concentrations ranging from 1 μM to 7.5 mM. These antisense sequences (**Table S1**) are complementary to the central domain of our DMRD, which also comprises the EDM. Hybridization of the antisense DNA to the DMRD hinders ATP-induced structure-switching of the aptamer by increasing separation between the two segments of the DDM. As a consequence, higher concentrations of ATP must be present to outcompete the effect of the antisense DNA, thereby shifting the dynamic range of the DDM for ATP detection.

**Figure 2.**
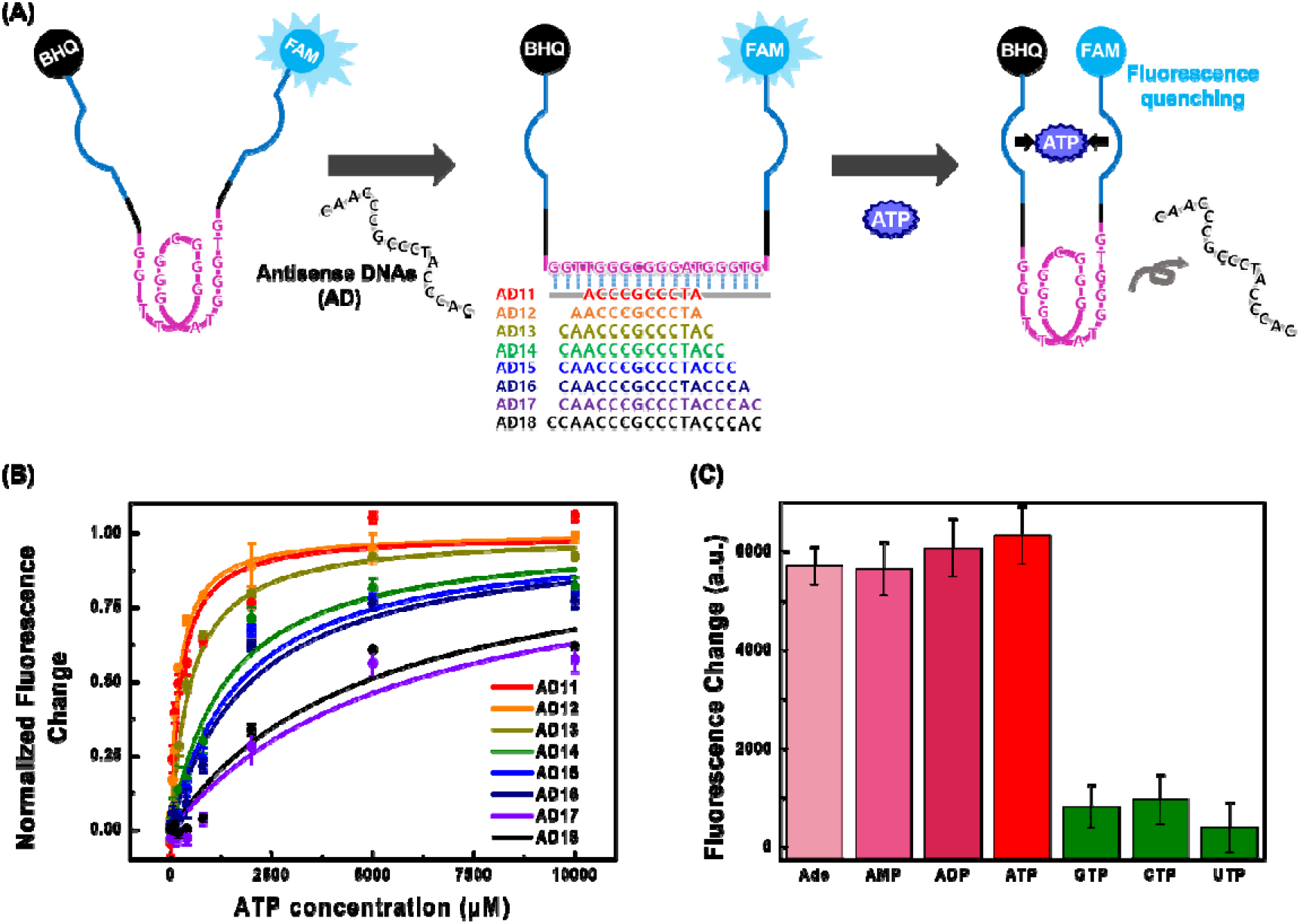
Tuning the ATP response of the DDM with antisense DNAs. (A) The target detection range of the DDM can be fine-tuned with antisense DNA strands that hybridize with the EDM. This hybridization hinders target-induced structure-switching, requiring higher ATP concentrations to generate fluorescence quenching. (B) Antisense strands with lengths ranging from 11–18 nt (AD11 to AD18) differentially affected the ATP response of this module, yielding DDMs that can cover a broader ATP detection range from 1–7,500 μM. (C) DDM target specificity in the presence of AD11. All ATP analogues were present at 1 mM. GTP, CTP and UTP failed to yield strong signal, whereas adenosine (Ade) and its derivatives (ATP, ADP, and AMP) produced a similar fluorescence change.

One can precisely tune the target detection range of the DDM by changing the length of the antisense DNA (**Figure 2B**). For example, a 13-nt antisense DNA produced a sensor with a half-maximal effective concentration (EC_50_) of 494 μM (**Table S2**), while the DDM without the antisense DNA yields an EC_50_ of 23.1 μM. As a result, the original target detection range of the DDM (1–100 μM) can be shifted to 57.2–2,998 μM (**Table S3**). With a shorter, 11-nt antisense DNA, we can lower this EC50 to 238 μM, achieving a detection range of 33.9–1,782 μM. In contrast, a much longer antisense strand (18 nt) produced a >20-fold greater EC50 value of 4,768 μM, such that our DDM can respond to a range of ATP concentrations from 346.8-7,440 μM.

As expected, the relationship between the length of the antisense DNA and the resulting EC50 value follows an Arrhenius plot (**Figure S1**). This is because the main interactions expected from our DDM—between the DMRD and the antisense DNA, and between the aptamer and ATP—are based on Gibbs free energy change (ΔG). From the extrapolation of our Arrhenius plot, we calculated the EC_50_ value for the antisense-free DDM, which is equivalent to the Kd of the ATP-binding aptamer (19.98 μM). Microscale thermophoresis measurements confirmed our calculations of the inherent binding affinity of the aptamer, yielding a K_d_ value of 23.1 μM based on Langmuirian binding between the target and the aptamer (**Figure S2**). The hybridization of antisense DNA interferes with the DDM’s structure-switching, allowing the binding behavior of the DDM to follow our hybridization energy-based Arrhenius plot, so that the DDM paired with an antisense strand exhibits a much higher EC50 than the Kd of the antisense-free DDM. Thus, DDM constructs coupled with varying lengths of antisense DNA can achieve finely-tuned binding curves that enable an ATP detection range spanning 1–7,500 μM (**Figure 2B**).

Our DDM consistently remained highly specific for ATP and other adenosine-derived molecules, without responding to other analytes with chemically similar structures (**Figure 2C** and **Figure S3**). For example, when the DDM was hybridized with the 11-nt antisense strand (AD11), it produced a similar reduction in fluorescence signal in response to adenosine, ATP, ADP, and AMP, but other nucleoside derivatives such as GTP, CTP, and UTP generated minimal signal change (**Figure 2C**). Importantly, this target specificity of the DDM does not differ in the absence of antisense strand (**Figure S3**), indicating that the antisense strand does not alter the nucleobase-specificity of the DDM. We therefore conclude that the presence of the antisense DNA only affects the thermodynamics of ATP binding, and thus the module’s target sensitivity and detection range.

### Optimizing EDM performance

We were likewise able to tune the responsiveness of the EDM by modulating the chemical conditions of the assay, which enabled us to achieve a dynamic range spanning roughly 1–500 mM with the same construct. Tris(hydroxymethyl)aminomethane (Tris) binds antagonistically to the EDM, thereby decreasing its catalytic activity, and can thus be used to shift its target detection range (**Figure 3A**). This Tris-dependent catalytic inhibition is a new finding, and the underlying mechanism may be related to a previous report that Tris can trap intermediate radicals (·OH) and thereby block the peroxidation of cellular lipids.^24–25^ In the absence of Tris, the catalytic activity of the EDM is dramatically enhanced by the presence of ATP, which serves as a cofactor for the DNAzyme-catalyzed peroxidase reaction. The presence of Tris molecules has an inhibitory effect and reduces the rate of this reaction, such that a higher concentration of ATP is needed to restore the DNAzyme’s peroxidase activity and thus generate an absorbance signal.

**Figure 3.**
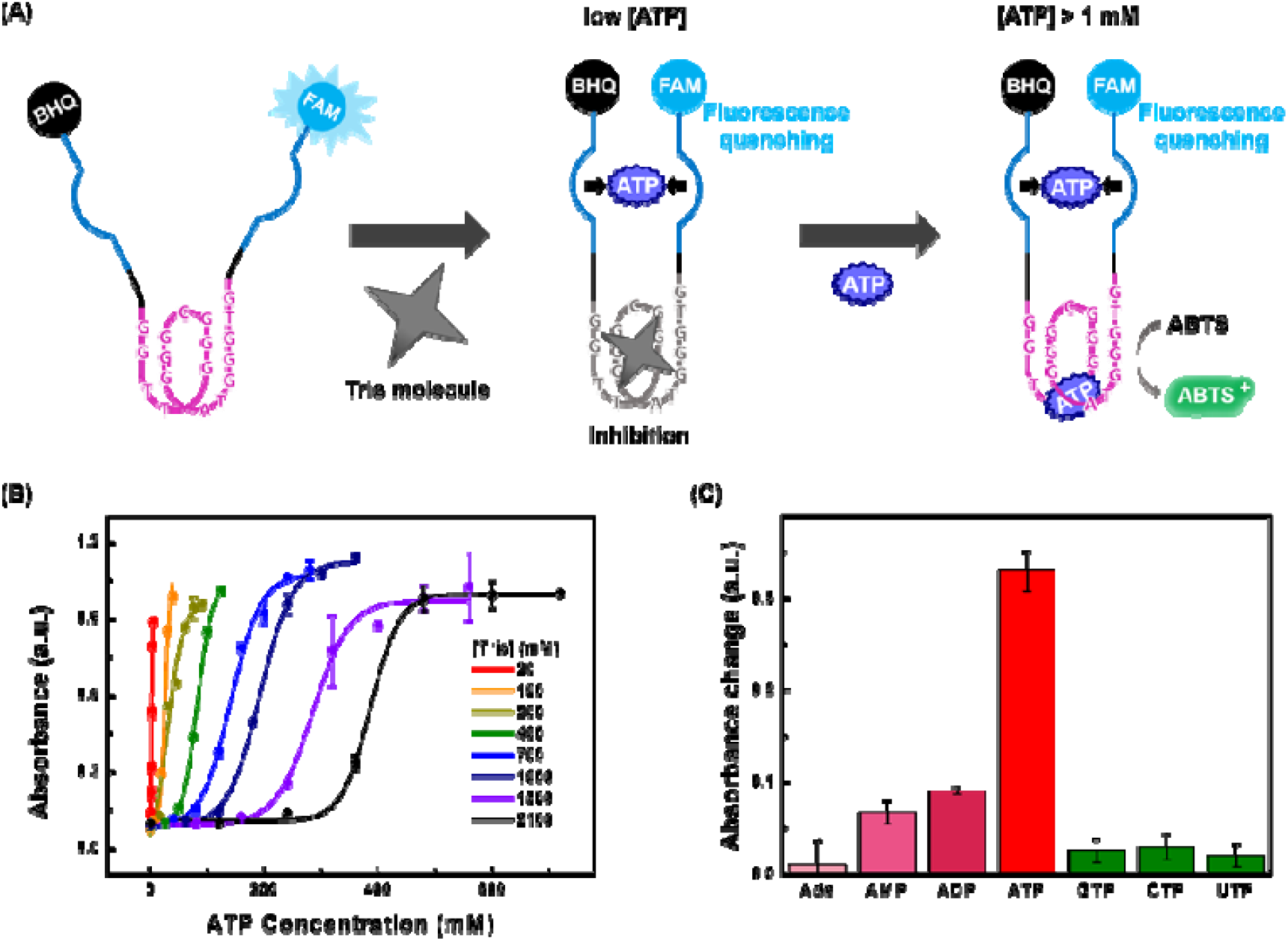
The EDM responds to ATP at millimolar concentrations, with a dynamic range that can be tuned by the presence of Tris. (A) Tris acts as a catalytic inhibitor of the EDM at low or moderate ATP concentrations, requiring more ATP to restore the reduced catalytic activity. (B) We obtained different calibration curves for ATP detection at different concentrations of Tris, where the ATP concentration required to generate a readout increased in proportion to the Tris concentration. With this tuning strategy, the EDM can achieve ATP detection at a range of ~1–500 mM. Each peroxidation reaction was performed at room temperature for 1 h. (C) Target specificity of our EDM in 20 mM Tris, with 3 mM of various ATP analogues. Adenosine, GTP, CTP, and UTP failed to generate a readout; ATP showed very strong catalytic enhancement, while ADP and AMP exhibited much weaker activity.

We experimentally confirmed our ability to finely tune the dynamic range of the EDM by adjusting the concentration of Tris from 20 mM to 2.1 M, obtaining eight different calibration curves for ATP detection (**Figure 3B**). At the lowest Tris concentration (20 mM), the EDM remained sensitive to relatively low ATP concentrations, ranging from 1.36–4.19 mM. At an intermediate concentration (700 mM Tris), the EDM was responsive to ATP concentrations ranging from 88.6–198 mM. Very high Tris concentrations greatly reduced this module’s sensitivity, and at 2.1 M Tris, the EDM exhibited a dynamic range that spanned sub-molar concentrations (282–506 mM). Thus, manipulating the chemical environment in this fashion can extend the ATP detection range achieved by the EDM from as low as 0.74 mM to as high as 506 mM.

Like the original DNAzyme,^22^ our EDM retains excellent specificity for ATP in the presence of Tris, and can even distinguish between ATP and other adenosine-based compounds (**Figure 3C**). ATP analogues such as GTP, CTP and UTP failed to generate an enhanced absorbance readout from the EDM in the presence of 20 mM Tris. ADP and AMP produced a much weaker response, while adenosine itself showed negligible influence on the peroxidation reaction. We identified the same ATP specificity in experiments without Tris (**Figure S4**), indicating that the addition of Tris does not affect the ATP specificity of the DNAzyme. Moreover, our observations indicate that unlike the DDM, our EDM can discriminate ATP from other adenosine-containing molecules based on the number of phosphate groups, and thus exhibits superior specificity compared to the DDM. This is because ATP plays a very specific role as a cofactor in the peroxidase activity of the EDM, where it activates H2O2 and provides the energy for electron transfer,^26^ whereas ATP binds to the DDM purely on the basis of intermolecular interactions.

### Dual readouts for ultra-broad detection range

We have demonstrated that these two modules, combined with the respective tuning mechanisms described above, enable the DMRD sensor to achieve an exceptionally broad dynamic range for ATP detection—from 1 μM to 500 mM, spanning more than five orders of magnitude. Consequently, we can quantitatively determine the ATP concentration in a sample by using this dual-readout system with two distinctive signals (**Figure 4**). For example, an ATP-containing sample can be added to microplate wells containing the DMRD with varying lengths of antisense strands or concentrations of Tris, and both fluorescence and absorbance readouts can be measured from each well. If the unknown sample has an ATP concentration within the dynamic range for detection in a particular condition, its concentration can be represented by the resulting specific fluorescence and absorbance intensities.

**Figure 4.**
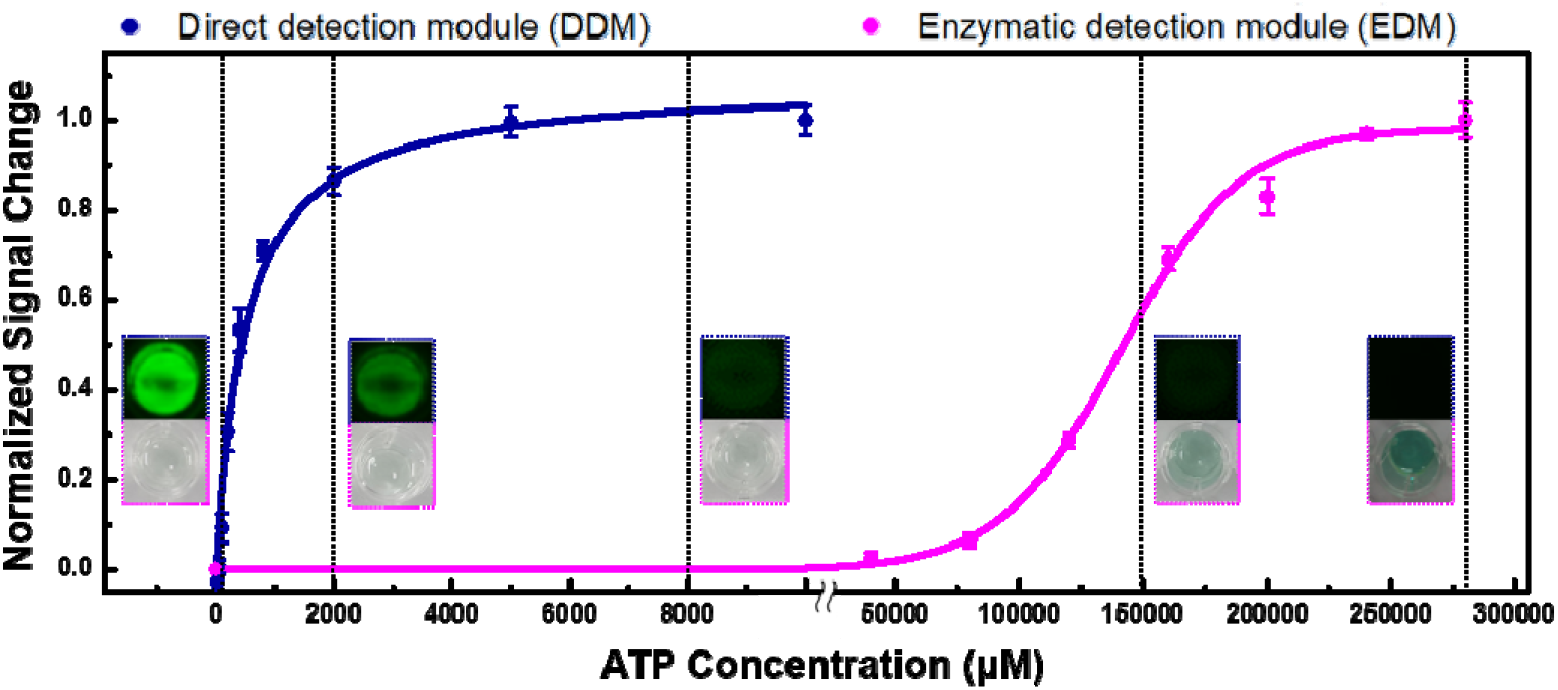
Applying the DMRD for ATP detection in different target concentration ranges. The concentration of ATP in an unknown sample can be represented by our dual readout (fluorescence and absorbance) platform in a 96-well microplate, where each well contains a different length of antisense DNA or Tris concentration.

## Conclusion

In this work, we integrated two different ATP-sensing modules in a single DNA molecule, which we have termed the DMRD, to achieve an extremely broad target detection range. The DDM employs a split aptamer coupled with a fluorophore/quencher pair to produce a binding-induced fluorescence change at micromolar ATP concentrations, where the dynamic range can be modulated through the introduction of complementary antisense strands of varying lengths. In parallel, the EDM incorporates a DNAzyme that employs ATP as a cofactor, generating an absorbance change at millimolar ATP concentrations, and the dynamic range of this module can be further tuned by manipulating the concentration of Tris in the reaction mixture. By integrating these two sensing modules, our *in vitro* ATP-sensing platform can achieve ATP detection at concentrations ranging from 1 μM to 500 mM. This spans a 500,000-fold concentration difference, which represents the widest sensing range achieved by a single DNA-based biosensor.

It is important to note that we do not have to generate variants of the bioreceptor with different detection ranges through complicated processes such as mutational screening. In addition, we can make use of alternate signaling modalities in our DMRD construct by substituting FAM or ABTS with different signaling substrates. For instance, if we swap ABTS with luminol, the DMRD can be converted to a two-color fluorescence signaling system. In conclusion, we have demonstrated here the use of two different detection mechanisms in a monolithic biosensor construct to achieve an extremely wide range of target detection. Although our current modular approach is limited by the availability of nucleozymes, we believe that ongoing advancements in nucleic acid library screening techniques^27–32^ will allow their accelerated discovery, making our strategy available for many other molecular targets.

## Acknowledgments

This work was supported by NRF(National Research Foundation of Korea) Grant funded by the Korean Government(NRF-2017R1C1B3012050 and NRF-2018-Fostering Core Leaders of the Future Basic Science Program/Global Ph.D. Fellowship Program).

