## Supplemental Information for "A Dual-sensing DNA Nanostructure with an Ultra-broad Detection Range"

**Table of Contents**

1. **Materials and Methods**
   1.1 Reagents and Materials
   1.2 Detection range measurement of the DMRD
   1.3 Tuning target detection range of the DDM
   1.4 Tuning target detection range of the EDM
2. **Supplementary Figures and Tables**

Table S1: DNA sequences

Table S2: ATP detection range of the DDM

Table S3: hybridization energies of antisense DNAs and EC_50_ value of the DDM

Figure S1: Arrhenius plot

Figure S2: Fitting graph for obtaining K_d_ value of the ATP aptamer

Figure S3: Antisense-free target specificity of the DDM

Table S4: ATP detection range of the EDM

Figure S4: Tris-free target specificity of the EDM

**1. Materials and Methods**

**1.1 Reagents and Materials**

DMRD and antisense DNAs AD11–18 were synthesized and purified by either Integrated DNA Technologies (Coralville, IA) or BIONEER (Daejeon, Korea). Adenosine triphosphate (ATP), Tris(hydroxymethyl)aminomethane (Tris), Tris hydrochloride, sodium chloride (NaCl), magnesium chloride (MgCl_2_), potassium chloride (KCl), Triton X-100, HEPES, and hydrogen peroxide (H_2_O_2_) were purchased from GA Biochem (Chuncheon, Korea). Adenosine diphosphate (ADP), adenosine monophosphate (AMP), adenosine, dimethyl sulfoxide (DMSO), sodium hydroxide (NaOH) and hemin were purchased from Sigma-Aldrich (St. Louis, MO). Guanosine triphosphate (GTP), cytosine triphosphate (CTP) and uracil triphosphate (UTP) were purchased from Tech and Innovation (Chuncheon, Korea). 2,2’-Azino-bis(3-ethylbenzothiazoline-6-sulphonic acid) (ABTS) was purchased from Roche (Basel, Switzerland).

**1.2 Detection range measurement of the DMRD**

We measured the affinity of the DDM for ATP via microscale thermophoresis (2bind GmbH; Regensburg, Germany). A titration series from 21.4 nM to 350 μM ATP was prepared in 1X ATP Binding Buffer (ATPBB; 20 mM Tris-HCl, 300 mM NaCl, and 5 mM MgCl_2_, pH 7.5), with the same concentration of DMRD (2 nM) added to each. The binding behavior of each sample was analyzed in the Monolith NT.115 pico device (NanoTemper Technologies GmbH; Munich, Germany). This experiment was performed in duplicate. In parallel, we also conducted FAM fluorescence visualization in 1.5 ml tubes containing 100 μl 1X ATPBB with 5 μM DMRD, in which we compared ATP-induced fluorescence decrease with 1 mM ATP or a negative control without ATP. Fluorescence signals were obtained using the Azure c600 (Azure Biosystems; Dublin, CA) at room temperature.

To confirm the ATP concentration range that can be detected by the EDM, we performed an ATP-assisted peroxidation test. Solutions containing 100 nM DMRD and 5 mM HEPES (pH 7.5) were prepared in 1X ATP Reaction Buffer (ATPRB; 10 mM KCl, 200 mM NaCl, and 0.002% (v/v) Triton X-100). To this solution, we added 5 mM H_2_O_2_ and 500 μM ABTS, and then incubated at 25 ℃ for 10 min. These reaction mixtures were challenged with varying concentrations of ATP (0–5 mM) and then transferred to a flat-bottomed transparent 96-well microplate (Greiner; [Kremsmünster](https://www.google.co.kr/search?newwindow=1&q=kremsmunster&stick=H4sIAAAAAAAAAOPgE-LSz9U3SEvONjGuUgKzjUxLCouMtLSyk63084vSE_MyqxJLMvPzUDhWGamJKYWliUUlqUXFi1h5sotSc4tzS_OKgXwAmA4nBlUAAAA&sa=X&ved=2ahUKEwjx06Ow2ezhAhVxw4sBHauoCJoQmxMoATAOegQICxAE), Austria), after which 200 nM hemin was added to each well (final reaction volume: 50 μl). The reactions were incubated at 25 ℃ for 1 h. Absorbance increases at 418 nm were measured with the Spark 10M microplate reader, with experiments performed in triplicate. In parallel, we performed naked eye visualization of ABTS^+^-associated color change (green) in 1.5 ml tubes, in which we compared ATP-assisted peroxidation in a 100 μl sample with 6 mM ATP versus a negative control without ATP, with results recorded by the digital camera of a Samsung Galaxy S9+ smartphone.

**1.3 Tuning target detection range of the DDM**

We prepared eight samples of 100 nM DMRD in 1X ATPBB, and then added 150 nM of an antisense DNA (AD11­–18) to each reaction (final volume: 100 μl). Samples were then heated to 95 ℃ for 10 min followed by ice-quenching for 10 min, and finally 10 min at room temperature. All samples were challenged with varying ATP concentrations (0–10,000 μM) and incubated for 30 min at 25 ℃, and then transferred to a flat-bottomed black 96-well microplate (Greiner; [Kremsmünster](https://www.google.co.kr/search?newwindow=1&q=kremsmunster&stick=H4sIAAAAAAAAAOPgE-LSz9U3SEvONjGuUgKzjUxLCouMtLSyk63084vSE_MyqxJLMvPzUDhWGamJKYWliUUlqUXFi1h5sotSc4tzS_OKgXwAmA4nBlUAAAA&sa=X&ved=2ahUKEwjx06Ow2ezhAhVxw4sBHauoCJoQmxMoATAOegQICxAE), Austria). Fluorescence signals were monitored by Spark 10M microplate reader with the following settings: excitation = 485 nm, emission = 535 nm, integration time = 40 μs. This experiment was performed in triplicate.

We tested DDM selectivity in the presence of AD11, with the reaction conditions and heat-treatment described above. The concentration of ATP, ADP, AMP, adenosine, GTP, CTP, or UTP was 1 mM. Fluorescence signals were monitored by Spark 10M microplate reader, with experiments performed in quadruplicate.

**1.4 Tuning target detection range of the EDM**

We prepared multiple sets of eight experimental samples (volume: 50 μl) containing 100 nM DMRD in 1X ATPRB with varying Tris concentrations (20 mM–2.1 M). After incubation at 25 ℃ for 10 min, each set of samples was challenged with varying ATP concentrations (1–720 mM), after which 200 nM hemin was added. The reaction was incubated at room temperature for 1 h. The increase of absorbance at 418 nm was measured by Spark 10M microplate reader, with experiments performed in triplicate.

We tested EDM selectivity in 20 mM Tris reaction conditions as described above, where each ATP analogue was present at 3 mM. Absorbance was monitored at 418 nm by Spark 10M microplate reader, with experiments performed in triplicate.

**2. Supplementary Figures and Tables**

**Table S1.** Sequences of DNAs used in this work. Blue sequences show components of the DDM; magenta sequences show the EDM.

| **Name** | **Sequence (5’ 🡪 3’)** |
| --- | --- |
| DMRD | FAM-AAT TCT GGG GGA GCC TTT TGT GGG TAG GGC GGG TTG GTT TTG CCC CGG AGG AGG AAT T-BHQ1 |
| AD11 | ACC CGC CCT A |
| AD12 | AAC CCG CCC TA |
| AD13 | CAA CCC GCC CTA C |
| AD14 | CAA CCC GCC CTA CC |
| AD15 | CAA CCC GCC CTA CCC |
| AD16 | CAA CCC GCC CTA CCC A |
| AD17 | CAA CCC GCC CTA CCC AC |
| AD18 | CAA CCC GCC CTA CCC AC |

**Table S2.** ATP detection range of the antisense-tuned DDM.

| **Antisense strand (# reflects length in nt)** | **Detection range (μM)** |
| --- | --- |
| AD11 | 33.9 ~ 1,782 |
| AD12 | 29.7 ~ 1,514 |
| AD13 | 57.2 ~ 2,998 |
| AD14 | 132.8 ~ 5,189 |
| AD15 | 160.9 ~ 5,678 |
| AD16 | 180.3 ~ 5,968 |
| AD17 | 389.3 ~ 7,663 |
| AD18 | 346.8 ~ 7,440 |

**Table S3.** The hybridization energies between antisense DNAs (AD11 to AD18) and the DMRD, and EC_50_ values in terms of ATP-induced structure-switching. The hybridization energies (△G, kcal/mole) between the antisense DNAs and the DMRD were calculated with the OligoAnalyzer Tool (Version 3.1) from Integrated DNA Technologies (IDT; Coralville, IA). Temperature was fixed at 298.15 K. EC_50_ values of our DDM increased depending on the length of antisense DNAs.

| **Name** | **Hybridization Energy (△G_hybridization_) (kcal/mole)** | **EC_50_ (μM)** |
| --- | --- | --- |
| AD11 | -24.87 | 238.23 |
| AD12 | -26.22 | 205.79 |
| AD13 | -28.17 | 494.17 |
| AD14 | -31.24 | 1,346.40 |
| AD15 | -34.31 | 1,694.10 |
| AD16 | -36.26 | 1,984.21 |
| AD17 | -37.6 | 4,827.45 |
| AD18 | -40.67 | 4,768.02 |

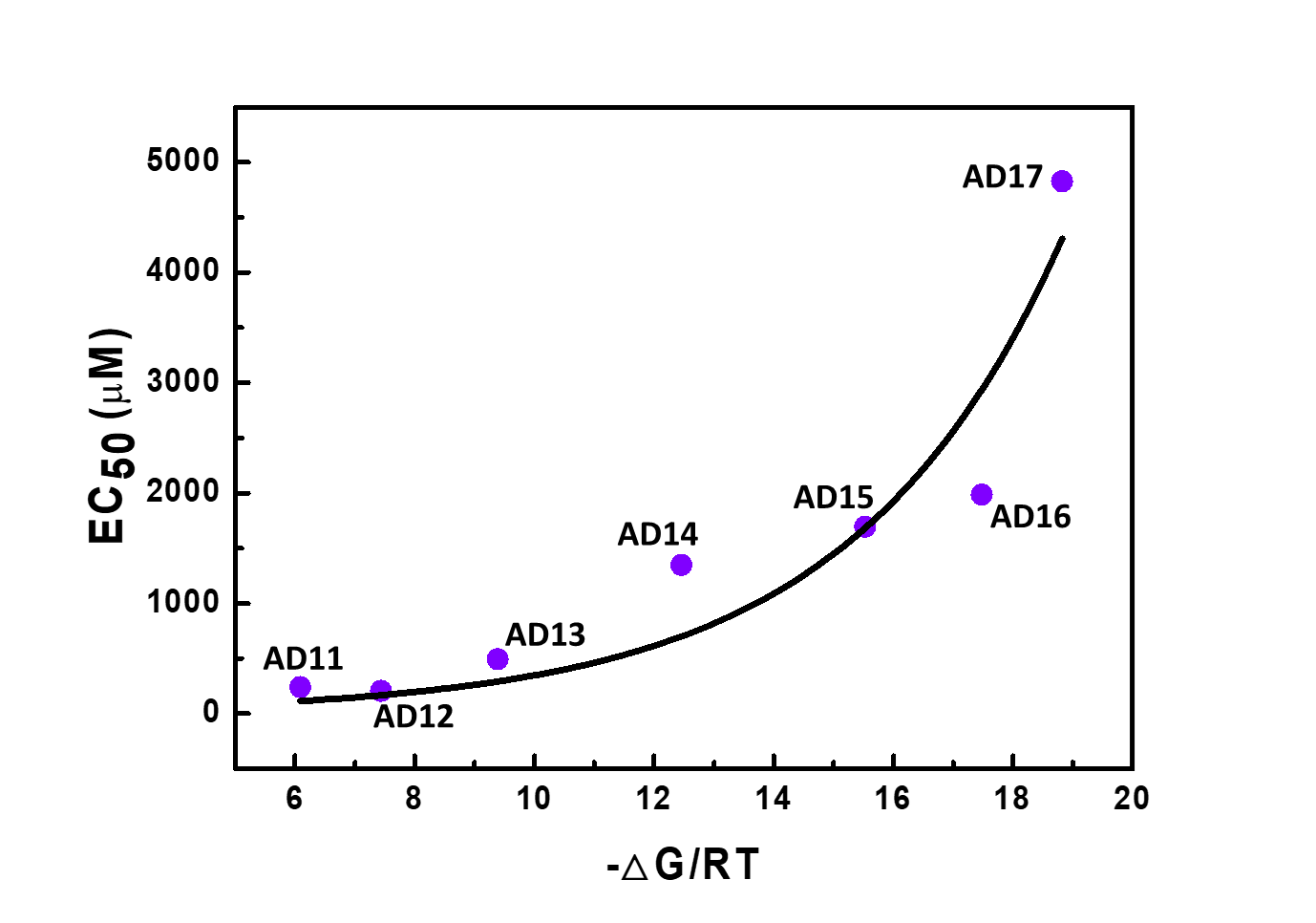

**Figure S1.** The increase of the EC_50_ value for antisense strands of increasing length relative to the Gibbs free energy changes (-△G/RT = (△G_hybridization_ - △G_mfold_)/RT) at room temperature (298.15 K). Regression, *y*: 19.98*exponential(0.285**x*); r^2^ = 0.873. △G_hybridization_ is the hybridization energy of antisense DNAs with the DMRD, and △G_mfold_ is the intrinsic folding energy of our DMRD (obtained by mfold software^1^). From extrapolation of this Arrhenius plot, we calculated a K_d_ value of 19.98 μM for the ATP-binding aptamer, which is quite similar to the experimentally measured value of 23.14 μM.

(1) Juker, M., Mfold web server for nucleic acid folding and hybridization prediction. *Nucleic Acids Res.* **2003**, *31*, 3406-3415.

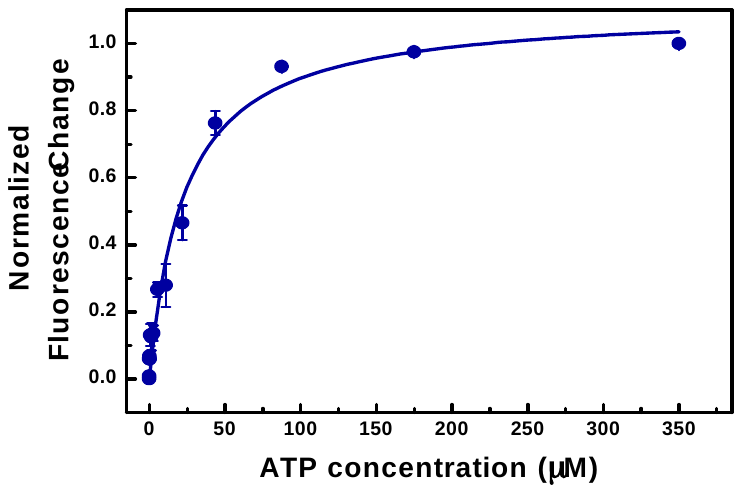

**Figure S2.** The fitting graph for obtaining the K_d_ of the ATP-binding aptamer based on microscale thermophoresis. Regression, y: 1.10**x*/(23.1 + *x*); r^2^ = 0.977.

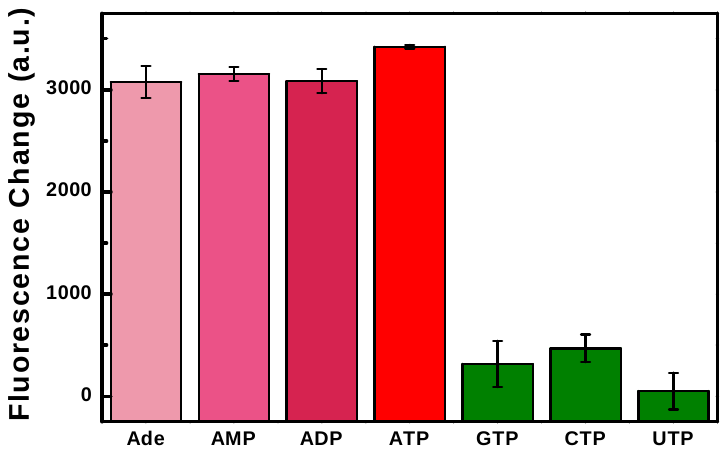

**Figure S3.** Target specificity of the antisense-free DDM. All ATP analogues, including adenosine (Ade), were present at 1 mM.

**Table S4.** ATP detection range of the EDM at varying Tris concentrations.

| **Tris concentration (mM)** | **Detection range (mM)** |
| --- | --- |
| 20 | 1.36 ~ 4.19 |
| 100 | 15.0 ~ 32.9 |
| 200 | 18.0 ~ 37.3 |
| 400 | 51.5 ~ 114 |
| 700 | 88.6 ~ 198 |
| 1000 | 126 ~ 273 |
| 1500 | 213 ~ 443 |
| 2100 | 282 ~ 506 |

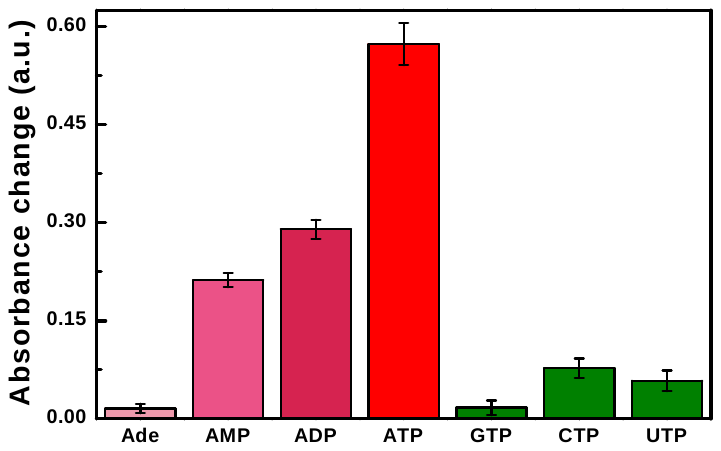

**Figure S4.** EDM specificity in the absence of Tris. All ATP analogues were present at 3 mM, and the peroxidation reaction occurred for 1 h.
